# Edge orientation perception during active touch

**DOI:** 10.1101/308759

**Authors:** Derek Olczak, Vaishnavi Sukumar, J. Andrew Pruszynski

## Abstract

Previous studies investigating the perceptual attributes of tactile edge-orientation processing have applied their stimuli to an immobilized fingertip. Here we tested the perceptual attributes of edge orientation processing when participants actively touched the stimulus. Our participants moved their finger over two pairs of edges–one pair parallel and the other nonparallel to varying degrees–and were asked to identify which of the two pairs was nonparallel. In addition to the psychophysical estimates of edge orientation acuity, we measured the speed at which participants moved their finger and the forces they exerted when moving their finger over the stimulus. We report four main findings. First, edge orientation acuity during active touch averaged 12.4°, similar to that previously reported during passive touch. Second, on average, participants moved their finger over the stimuli at 23.9 mm/s and exerted contact forces of 0.47 N. Third, across participants, there was no clear relationship between how people moved their finger or how they pressed on the stimulus and their edge orientation acuity. However, within participants, there appeared to be a weak effect of speed on performance whereby error trials included slightly faster finger movements; no equivalent effect was found for contact force. Fourth, consistent with previous work testing spatial acuity, we found significant correlation between fingertip size and orientation acuity such that people with smaller fingertips tended to have better orientation acuity.

## Introduction

Fine manipulation tasks–like buttoning a shirt or picking up a grain of rice–involve accurately determining an object’s orientation relative to the fingertips based on information arising from mechanoreceptors in the glabrous skin. Given the behavioral utility of edge orientation information, many studies have investigated the perceptual attributes of tactile edge orientation processing; almost all of this work applied edges to an immobilized finger and has consistently reported edge orientation acuity of ~15° (Bensmaia et al., 2008; Lechelt, 1992; Peters et al., 2015). We recently investigated edge orientation processing while participants actively touched and controlled an object’s orientation based only on tactile information (Pruszynski et al., 2018) and reported that edge orientation acuity expressed by the motor system in this context is ~3°, many times better than that measured via passive perceptual assays.

Here we examined the perceptual attributes of edge orientation processing during active movement. In our experiment, participants actively moved their finger over two pairs of oriented edges and identified which of the two edge pairs was not parallel by pushing a button next to that edge pair. We estimated tactile edge orientation acuity by titrating the orientation difference for the non-parallel edge pair. In addition to the psychophysical measures, because the edge pairs were mounted on six axis force transducers we could characterize the kinematics and dynamics of finger movement while performing the task.

We had three key aims. First, we wanted to characterize perceptual edge orientation acuity during active touch. Doing so would provide some insight into whether the difference in tactile edge orientation acuity between passive perceptual work and active object manipulation simply reflects active movement. If actively engaging in movement is, in and of itself, the source of better orientation acuity then tactile edge orientation acuity in our active perceptual task should be substantially better than previously reported in passive perceptual studies, perhaps even approaching the level we observed during object manipulation. Second, we wanted to quantify how people move and contact surfaces when perceptually discriminating oriented edges. Although such information is important for understanding the neural basis of tactile processing, what information exists in the context of fine spatial features is limited largely to the identification of characters from the roman alphabet, which feature a relatively crude assortment of edge orientations (Vega-Bermudez et al., 1991). Third, we wanted to test whether there exists a relationship between fingertip size and edge-orientation acuity during active movement. Several studies have reported that, on average, people with smaller fingertips perceive finer spatial details of touched surfaces than those with larger fingertips (Goldreich and Kanics, 2003, 2006; Van Boven et al., 2000), a difference thought to reflect the increased innervation density of Merkel and Meissner mechanoreceptors in smaller fingertips (Peters et al., 2009). To date, support for this idea comes from passively applied grating stimuli and it is unknown whether it generalizes to other tactile attributes or active touch. If the mechanistic explanation about innervation density is correct then tactile acuity in our active edge orientation task should also be negatively correlated with fingertip size.

## Methods

### Participants

A total of 91 university-aged adults volunteered for the study (37 male, 65 female; age range = 18 - 30). All participants were paid for their participation ($10) and reported having no known injuries or health conditions that affected their hand function or tactile acuity. Participants were asked about their sex but, as no detailed guidance was provided, we assume that their response related to how they identify (i.e. their gender). The study was approved by the Western University Humans Research Ethics Board and was performed in accordance with the Declaration of Helsinki.

### Apparatus

Participants placed their finger into a slot and sequentially moved their finger approximately 4 cm over two plates (Fig. 1A). Each plate was firmly attached to a six-axis force transducer (ATI Nano-17, Apex, NC) that provided information about the contact forces and torques generated by the participant. As in our previous work (Pruszynski and Johansson, 2014; Pruszynski et al., 2018), the plates, which were changed between trials, were covered with a nylon material (Toyobo EF-70-GB) embossed with two raised edges - 5 mm high, 5 mm wide at the top and 8 mm wide at the base).

**Figure 1.**
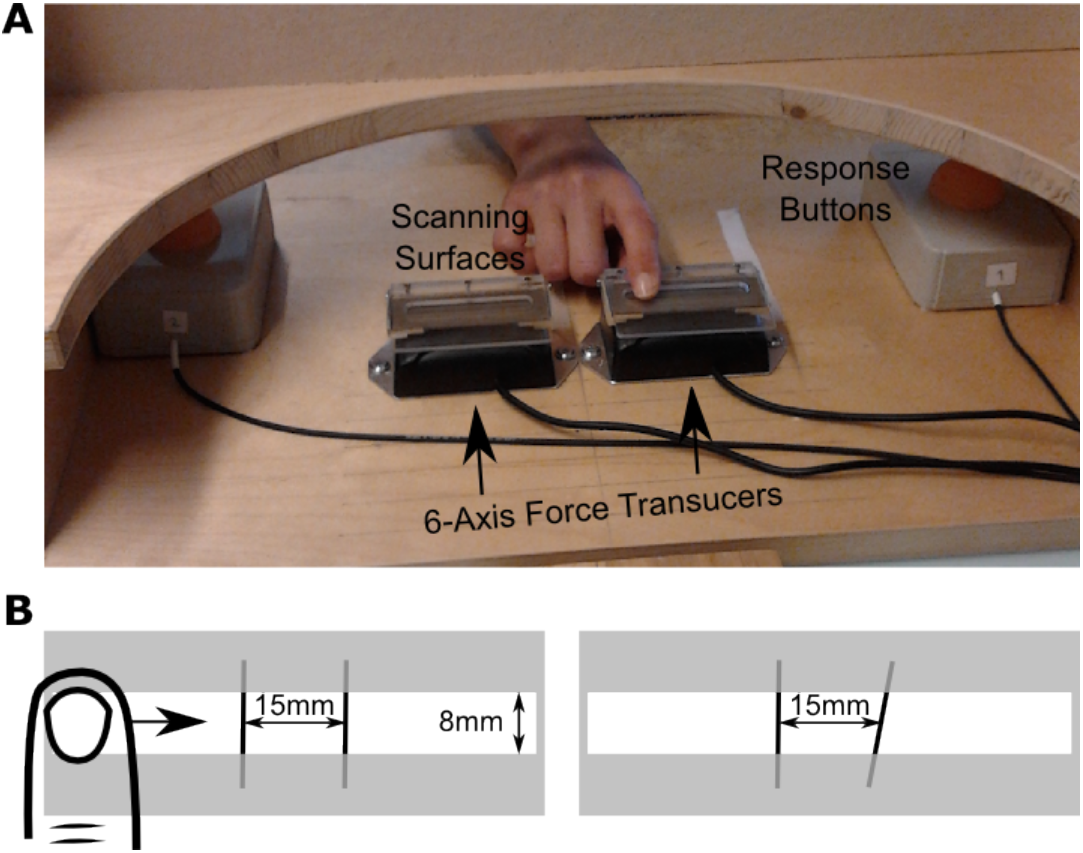
Experimental setup. (A) Photograph of one version of the experimental setup. Participants had their vision occluded and touched the tactile stimuli with their right index finger. The stimuli were mounted on six-axis force transducers which provided information about their contact forces were used to estimate their finger kinematics (see Methods). (B) An illustration of two experimental stimuli, one with parallel edges (left) and the other with non-parallel edges (right). Note that participants could only place their finger into the depicted slot (white area) such that finger movements were physically limited to one dimension.

The edge pairs on each plate could either be parallel or non-parallel. The parallel edges were oriented perpendicular to the direction of finger movement (Fig. 1B). We randomly used ten different parallel edge plates to ensure participants could not base their decision on some idiosyncratic imperfection in the parallel stimulus. The non-parallel edges included one edge perpendicular to the direction of finger movement and another edge rotated about its center by ±1-20° in 1° increments. We randomly drew from positive and negative edge orientation differences so that participants could not predict the overall geometry of the plates. For both parallel and non-parallel plates, the center of the plate corresponded to the midpoint between the centers of the edges and the center-to-center spacing between edges was 15 mm.

### Experimental procedure

The experimental task was to determine which of the two plates included non-parallel edges. We implemented a two alternative forced choice (2AFC) design. On each trial, a parallel plate and a non-parallel plate were presented to the participant randomly in one of two locations. The participant swiped their right pointer finger a single time across the left and then right plate and then pressed one of two buttons placed near each plate to indicate their judgement about which of the two plates contained the non-parallel edges. The only explicit instruction given to the participant about how to swipe was that they could only swipe in one direction (left-right or away-towards) without stopping and that they could not lift their finger. Participants had no difficulty meeting these requirements.

Stimulus progression followed a one-down ten-up progression, starting at 20°. That is, participants started with the non-parallel plate that contained edges with a 20° angular difference. If they correctly identified the non-parallel plate, they would move to the 19° edge and so on through the set of plates. When they answered incorrectly, the angular difference increased by 10 degrees to a maximum angular difference of 20°. In practice, this approach meant that participants almost always started a new run of trials with the 20° plate. Each participant performed four runs for each plate configuration block (see below) and we took the average of the minimum angular difference achieved in these four runs as a measure of their edge orientation acuity. Note that the first five trials in each plate orientation block were treated as practice and thus errors in the first five trials were not counted towards a participant’s runs.

Each participant performed two plate configuration blocks, where the stimuli were oriented left/right or away/towards relative to their body. This was done by keeping the plates in the same location but rotating them about their midpoint. For both configurations, the participant maintained the same hand orientation such that the direction of movement was aligned with the short axis of the finger pad for the horizontal swipe and the long axis of the finger pad for the vertical swipe. The assignment of which block the participant completed first was randomized and counterbalanced across participants.

After the psychometric task, we measured each participant’s finger pad area. Area measurements were performed by having the participant apply their right pointing finger to an ink pad and then to a slip of paper on the force transducer while visual and auditory cues guided them to one of two desired vertical force levels. We used force levels of 0.4 N and 0.8 N as they fell within the range observed in pilot studies and were consistent with previous studies examining forces during fine haptic exploration for perceptual report (Vega-Bermudez, Johnson & Hsiao, 1991). Slips were then placed on a forensic ruler and imaged at a fixed focal distance and the same lighting. Images were processed using standard contour select and area measurement tools within ImageJ (v1.50b, NIH, Bethesda, MD) with manual calibration of scale and sensitivity.

### Data acquisition and analysis

Custom software was created for data acquisition and processing (MATLAB R2015b; MathWorks Inc., Natick, MA). Data was acquired from the force transducers via a serial port with precise time stamps at a sampling rate of ~60Hz. The data was filtered with a low pass Butterworth filter (second order, two pass, 20Hz cut-off).

Center of pressure was calculated as the ratio of the torque caused by moving the finger along the slot (Ty) and the vertical contact force (i.e. the pressing force, Fz). For behavioral analyses, we focused on data ±2 cm from the midpoint of each stimulus plate. Average forces were based on all samples within this measure measurement window. Average swipe velocity was calculated by noting how long it took the finger to move over the 4 cm measurement window. Fingertip area was calculated as the average of the two force levels for each participant.

Two participants did not fully complete the experiment because of technical difficulties. Participant level averages for these participants is based on less than four runs in one of the two configurations. One participant did not yield usable data for finger speed and force data and is excluded from this analyses. One participant did not yield usable fingertip area data and is excluded from this analysis.

Statistical analyses were performed in MATLAB or JASP (Version 0.8.5.1, University of Amsterdam, The Netherlands). Descriptive values are presented as mean ± standard deviation (SD) unless otherwise noted. Statistical results were deemed significant if p < 0.05.

## Results

### Psychophysics

Participants had little difficulty performing the task and typically completed the full experiment in ~45 minutes in which time they performed 68 ± 20 trials.

Perceptual orientation acuity during our active touch experiment resembled that established in previous passive touch studies (Figure 2A). On average, our participants were able to reliably discriminate edges that differed by 12.4 ± 2.9° (median = 12.5°, 25-75th percentile = 10-14°). Figure 2B illustrates edge orientation acuity as a function of plate configuration. We found that participants showed significantly better orientation acuity (paired t-test, t_90_ = 6.2, p < 1.02 × 10^−7^, Cohen’s d = 0.65) for the away/towards orientation (11.2 ± 4.0°; median = 13.75, 25th-75th percentile = 11.25-15.5) as compared to the left/right orientation (13.6 ± 2.8°; Median = 10.75, 25th-75th percentile = 8.25-14.25). Across participants, there was a statistically significant correlation between orientation acuity for the two plate configurations (Pearson’s r = 0.44; p < 1.2 × 10^−5^).

We found no clear evidence that learning or fatigue influenced edge orientation acuity. There was no significant difference in orientation acuity for either the left/right (independent samples t-test, t_89_ = −0.38, p = 0.71, Cohen’s d = −0.080) or away/towards configuration (t_89_ = −0.07, p = 0.95, Cohen’s d = −0.014) as a function of whether it was done first or second in the experimental session. Nor was there a significant difference in orientation acuity between the 1st and 4th runs for either configuration (paired t-test, left/right: t_90_ = −1.4, p = 0.16, Cohen’s d = −0.15, away/towards: t_88_ = −0.87, p = 0.39, Cohen’s d = −0.093).

**Figure 2.**
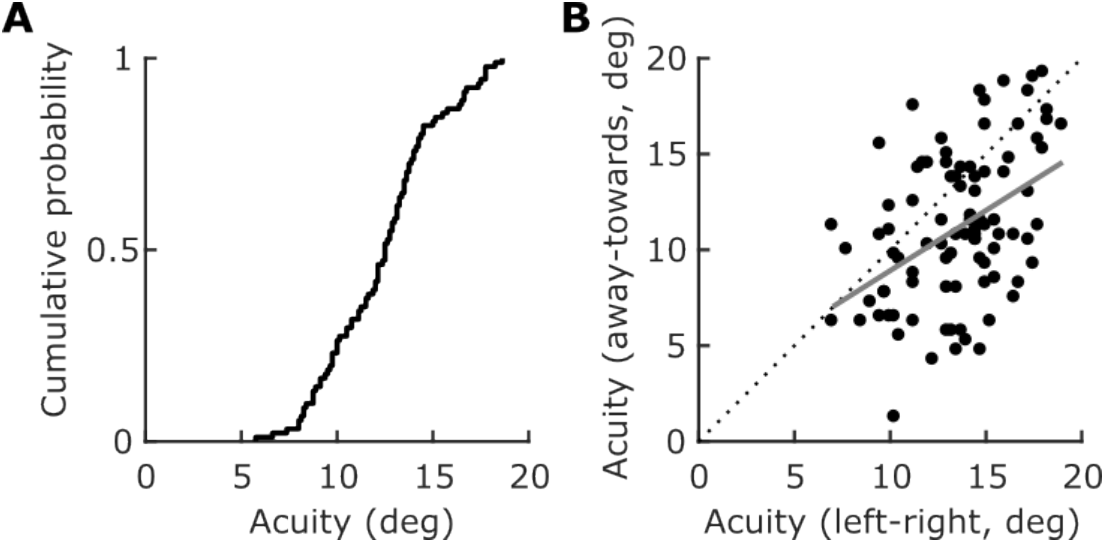
Orientation acuity. (A) Cumulative histogram showing average orientation acuity (across both plate orientations) in our sample. (B) Scatter plot relating orientation acuity for the left-right and away-towards plate orientations. Each data point represents an individual participant. The dashed line is the unity line. The gray line represents the outcome of a linear regression relating these variables.

### Finger movement and contact forces

Figure 3A illustrates how a representative participant moved their finger over the stimuli. As was typically the case, this participant moved at a relatively constant speed along the swiping direction in any given trial and did so in a relatively consistent manner across trials; average swiping speeds for this participant ranged from 20 to 45 mm/s. Figure 3B illustrates the downward contact forces exerted by the participant. As was typically the case, this participant exerted relatively consistent downward force within a trial but was much more idiosyncratic between trials, ranging from 0.1 to 0.7 N.

**Figure 3.**
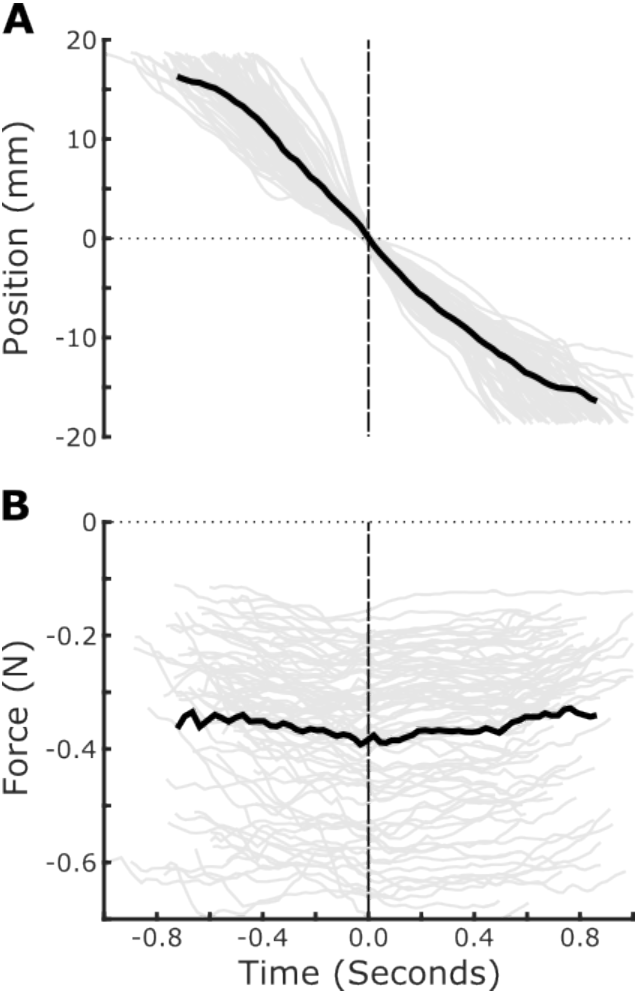
Kinematics and forces of a representative participant. (A) Finger position for a for a single participant (gray lines = single trials; black line = mean). The horizontal axis represents time and the vertical axis represents position relative to the center of the plate. Data is aligned to the time when participants reach the middle of the plate and, as such, all traces pass through the origin. (B) Same format as (A) but for finger forces whereby negative values indicate pressing on the plate.

Figure 4A and B summarize the finger speed and contact forces used by our participants. On average, participants moved their finger over the stimuli with a speed of 23.9 ± 15.4 mm/s and exerted a contact force of 0.47 ± 0.43 N. We found no reliable correlation between finger speed and contact force (Spearman rho = 0.025, p = 0.81). We also found no reliable difference in finger speed or contact force as a function of plate configuration (paired t-test, speed: t_89_ = 0.006, p = 0.995; force: t_89_ = 0.78, p = 0.43). Across participants, both finger speed (Pearson’s r = 0.78, p < 10^−6^; Fig. 4C) and contact force (r = 0.81, p < 10^−6^; Fig. 4D) were strongly correlated across the two plate configurations.

**Figure 4.**
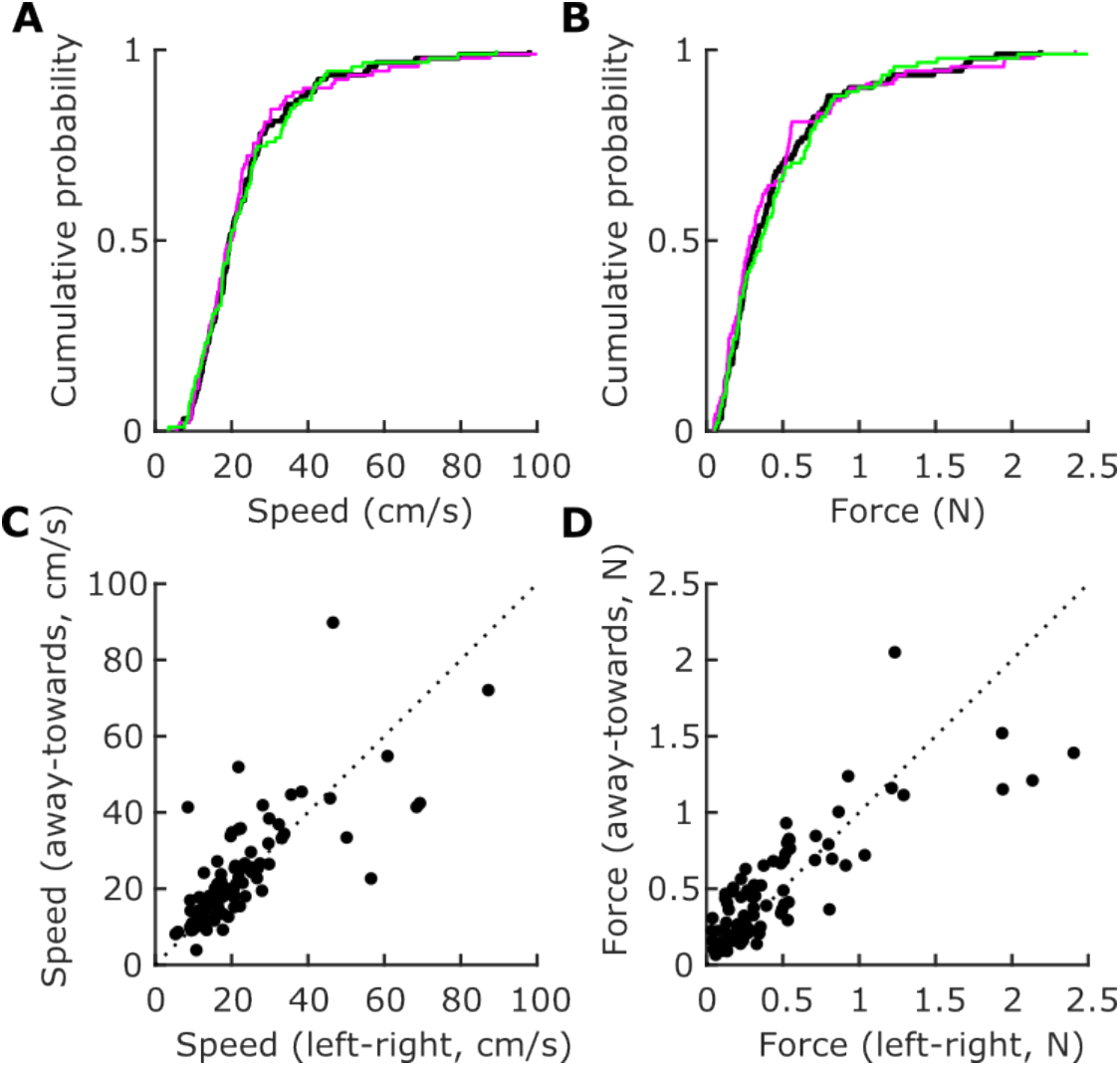
Kinematics and forces across the population. (A) Cumulative histogram of average finger speed for each participant (black line = mean of all trials; magenta line = mean for only left/right; green line = mean for only away/towards). (C) Scatter plot relating the finger speeds used by each participant in the left/right and away/towards plate configurations. (B,D) Same format as (A,B) but for finger forces.

We assessed whether the range of finger speeds and contact forces used by the different participants influenced edge orientation acuity. That is, did participants who expressed a particular movement pattern show better orientation acuity? This did not appear to be the case as we found no reliable relationship between how quickly a participant moved and how they ranked in terms of edge orientation acuity averaged across plate configurations (Spearman’s rho = −0.027, p = 0.80; Fig. 5A). We also found no reliable relationship between contact force and performance (rho = −0.121, p = 0.26; Fig. 5B).

**Figure 5.**
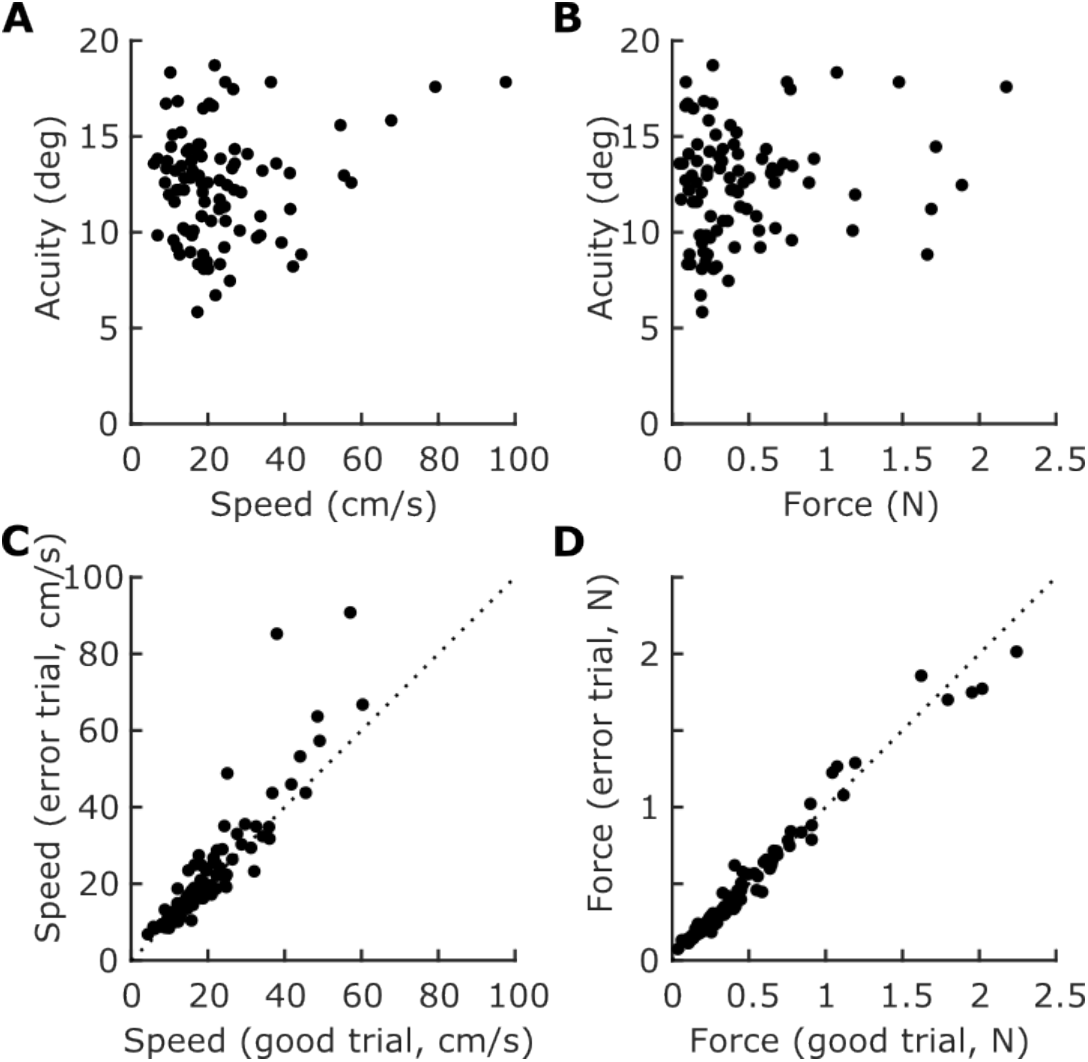
Movement parameters and performance. (A) Scatter plot relating average finger speed and orientation acuity for each participant (averaged across plate configurations). (B) Same format as (A) but for finger forces. (C) Scatter plot relating the average finger speed on error trials and the previous correct trials. Each dot represents a single participant. (D) Same format as (C) but for finger force.

We then examined the relationship between finger movement parameters and orientation acuity at the single participant level. We leveraged the fact that individuals used a range of speeds and contact forces to examine whether there was any trial-by-trial relationship between movement or force parameters and correctly identifying the non-parallel edge pair. Specifically, we analyzed whether last correct trial and the subsequent incorrect trial that ended each run showed reliable differences in speed and contact force. Here, we found a weak but reliable effect of speed (paired t-test, t_89_ = 2.11, p = 0.038) whereby participants moved slightly faster on the final error trial (22.9 ± 15.7 mm/s) than the last correct trial (21.3 ± 11.3 mm/s; Fig. 5C). Given the small effect, we also checked whether any reliable relationship existed between the second last and last correct trials and found no difference in finger speeds between these two correct trials (paired t-test, t_89_ = −0.96, p = 0.34). We found no reliable difference in contact force between the last correct trial and the error trial (previous trial: 0.49 ± 0.44N; final trial: 0.48 ± 0.43N; t_89_ = 1.46, p = 0.15; Fig. 5D).

### Effect of gender and fingertip size

Our results in active touch are consistent with previous work linking fingertip size to tactile acuity in passive situations (Peters et al., 2009). Figure 6A illustrates edge orientation acuity (averaged across plate configuration) as a function of self-reported gender. We found no reliable effect of gender on edge orientation acuity (t_89_ = 1.2, p = 0.245) though we did find a clear effect of gender on fingertip size (t_88_ = 4.75, p = 7.9 × 10^−6^; Fig. 6B). More notably, as in passive situations, there was a reliable correlation between fingertip size and edge orientation acuity (Pearson r = 0.24, p = 0.023; Fig. 6C) whereby people with smaller fingertips tended to discern more similar edge orientations.

**Figure 6.**
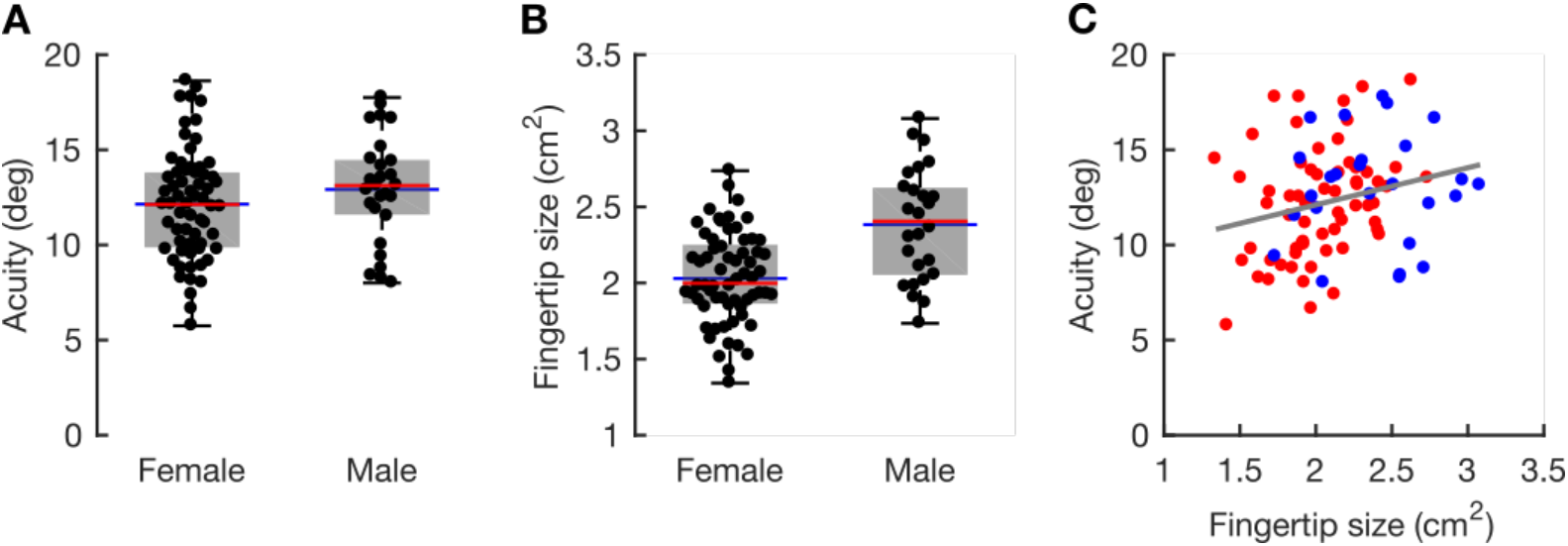
Effect of gender and fingertip size on behavior. (A) Box plot showing edge-orientation acuity as a function of self-identified gender. Each data point represents a single participant. Horizontal lines represent mean (blue) and median (red); the extent of the box represents the 25th and 75th percentile. (B) Same format as (A) but for fingertip size. (C) Scatter plot relating fingertip size and gender. The red and blue dots denote females and males, respectively. The line represents the linear fit collapsed across all participants.

We also tested whether there was any effect of gender or fingertip size on finger movements and contact forces. We found no reliable differences for either variable (speed: t_89_ = 1.39 p = 0.17, Cohen’s d = 0.319; force: t_89_ = 0.40 p = 0.70, Cohen’s d = 0.090). That is, men and women moved their fingers at similar speeds (men: 20.5 ± 12.2 mm/s; women: 25.3 ± 16.4 mm/s; Fig. 7A) and used similar contact forces (men: 0.50 ± 0.44 N; women: 0.46 ± 0.43 N; Fig. 7B). We also found no reliable correlation between fingertip size and finger speed (Pearson’s r = −0.056, p = 0.60; Fig. 7C) or contact force (Pearson’s r = −0.105, p = 0.32; Fig. 7D).

**Figure 7.**
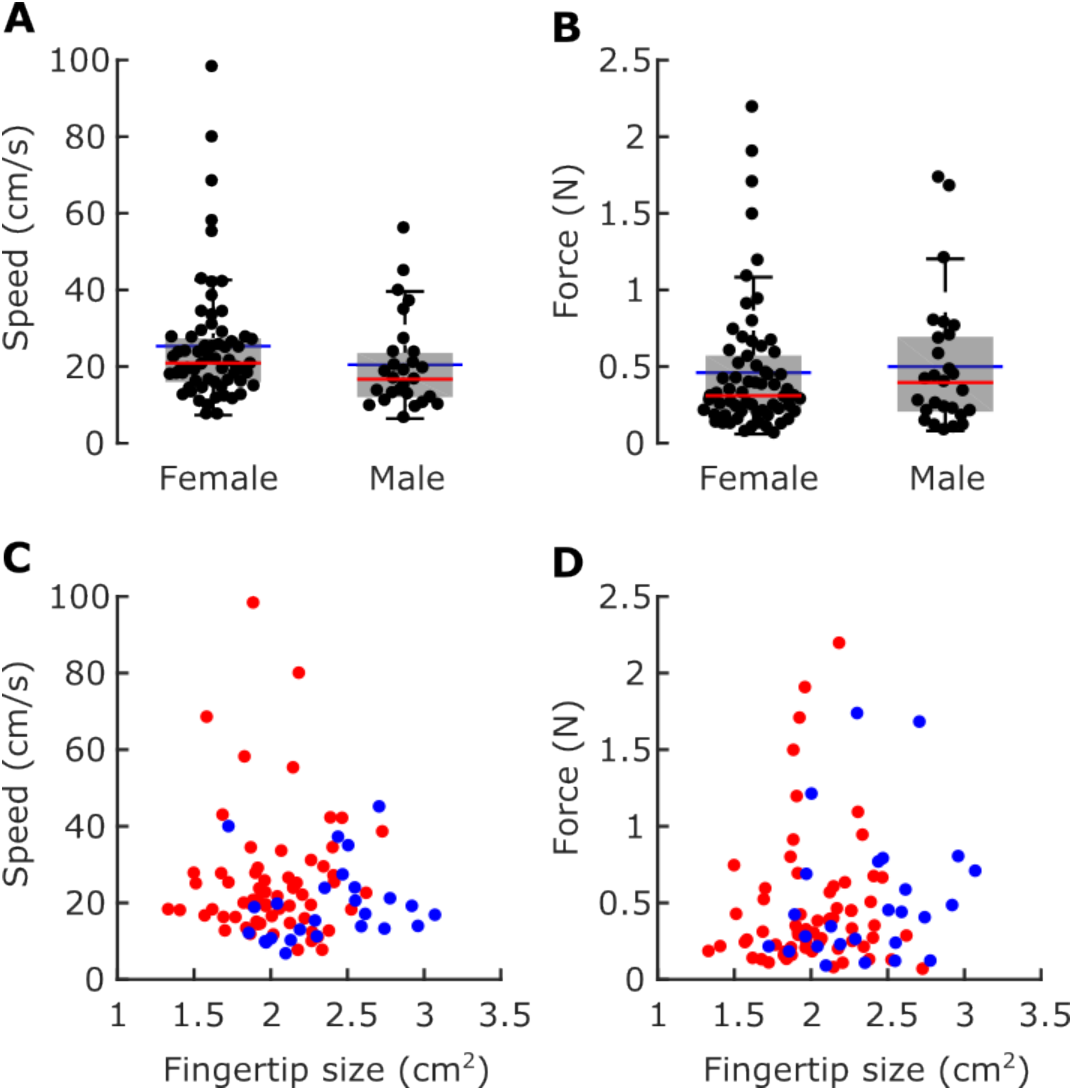
Movement parameters as a function of gender. (A,B) Same format as Figure 6A but for average finger speed and force, respectively. (C,D) Same format as Figure 6C but for average finger speed and contact force, respectively.

## Discussion

Our simple experiment yielded four main results. First, we found that edge orientation acuity during active touch, as measured in a psychophysical task, is similar to the many other perceptual studies where an immobilized finger is passively stimulated. Second, on average, we found that participants moved their finger over the stimuli at 23.9 mm/s and exerted contact forces of 0.47 N. Third, we found that, across participants, there was no reliable relationship between how people moved their finger or how they pressed on the stimulus and their edge orientation acuity. However, within participants, there was a weak but reliable relationship between finger speed and orientation acuity: trials where participants failed to discriminate edge orientation involved slightly faster finger speeds than the previous trial where they succeeded. Fourth, consistent with previous work looking at the limits of tactile spatial acuity, we found a correlation between fingertip size and edge-orientation processing such that people with smaller fingertips tended to show better acuity. Below we briefly discuss our main findings.

### Active touch versus passive touch

Tactile orientation acuity has typically been studied by stimulating an immobilized patch of skin and having the participant perceptually discriminate or identify edge orientation. A range of studies indicate that, for edges spanning a large part of the fingertip, perceptual sensitivity to edge orientation during passive touch is between 10° and 25° (Bensmaia et al., 2008; Lechelt, 1992; Peters et al., 2015). This finding appears robust across laboratories and when manipulating a host of stimulation parameters and psychophysical assays in the same basic experimental setup (Bensmaia et al., 2008). In a recent paper, we took a different approach by assessing edge orientation acuity in the context of object manipulation (Pruszynski et al., 2018). Rather than explicitly report their edge orientation perception, we had participants actively touch a pointer and align it to a preset configuration. Since the participants could not see the pointer, the only cue they had about its initial orientation was a raised edge on the contact surface that they touched when performing the action. Participants performed this task without difficulty, and were able to express edge orientation acuity of ~3° for edges spanning their entire finger, substantially better than during previous perceptual studies.

Here we assessed whether the orientation acuity advantage during object manipulation could be attributed to the act of actively generating a movement. This seems a reasonable possibility as active movement could, in principle, allow participants to optimize when and how the object was contacted, and thus increase the amount of edge orientation information being extracted. However, we found that people actively moving their finger to perceptually discriminate edge orientations showed tactile orientation acuity far from what they can show in the context of object manipulation - suggesting that such acuity does not arise because of movement per se. Indeed, we suspect that the acuity benefit relates to the specific demands of grasping and manipulation, where sensory signals arising from the skin contribute to the online control of the hand and digits (Johansson and Flanagan, 2009; Pruszynski and Johansson, 2014; Pruszynski et al., 2016, 2018).

Edge orientation acuity in the present task was, on average, within the range of previous passive tactile studies. This finding is consistent with previous work showing that people do not gain a significant perceptual advantage in terms of fine spatial acuity, and other tactile cues such as roughness, by actively moving their finger (Heller, 1989; Lamb, 1983; Lederman, 1981; Verrillo et al., 1999). For example, people are equally able to identify roman characters and show the same patterns of errors when they themselves move their finger relative to the stimulus and when the finger is stationary and the stimulus is moved by the experimenter (Vega-Bermudez et al., 1991). This is not to say that active touch does not provide benefits. There are many tasks where active movement yields better performance though these are typically in the context of haptic exploration of objects rather than fine spatial features of touched objects like edge orientation (Chapman, 1994; Gibson, 1962; Lederman and Klatzky, 1987).

It is important to note that we did not explicitly compare passive and active touch in our experiment so it is possible that actively moving improved performance, just not to the level we found in our object manipulation study. And the potential edge orientation acuity during active may have not reached our previously reported levels because of differences in the precise experimental protocol and thus available sensory information. For example, the biggest difference between this task and our previous motor control task is that the former included tangential motion between the skin and the stimulus whereas the latter was largely indentation. Although we would expect that the presence of tangential motion would, if anything, favor the current experiment in terms of available information, we did not directly test this idea. Another limitation in this regard is that we chose an experimental approach that did not force people to their perceptual limit - that is, we implemented a relatively fast protocol rather than an extensive search for perceptual threshold. It seems inevitable that such an approach would lower our estimate of perceptual threshold (Wong et al., 2013), as would providing people much more practice and experience with the task. Nevertheless, given our experience thus far, it seems very unlikely that they would increase their performance four fold to the levels we readily saw in motor control where participants had a similar level of experience and practice as in the present experiment.

### Haptic strategies for fine form discrimination

We wanted to address how participants move their finger when performing perceptual edge orientation discrimination. Similar to previous work in the context of tactile letter identification we found that, on average, participants moved their finger at ~20 mm/s (Vega-Bermudez et al., 1991). Although there was variability across participants, each individual showed relatively consistent speed between trials. Contact forces were, on average, ~0.5 N, but were variable both across participants and across trials for a given participant.

Although we did not actively manipulate speed, we found no robust relationship between how quickly a participant moved their finger and their edge orientation acuity. There are several reasonable interpretations of this finding. First, it may be that participants adjust their speed in a way that optimizes orientation information extraction given their particular finger geometry, skin mechanics and peripheral neural organization and other factors. Second, participants may simply use some habitual speed that is unrelated in any meaningful way to the task because how they move has relatively little effect on orientation acuity. Third, it may be that participants did not have sufficient time to discover an optimal finger speed given that we designed our experiment to provide minimal training and learning. An interesting follow-up would be to provide extensive training and test whether under more extreme conditions participants would optimize their behavior and gain a perceptual advantage in the process.

### Fingertip size and orientation acuity

We did not find a reliable effect of gender on edge orientation acuity. However, consistent with previous work investigating spatial tactile acuity with passive stimulation (Peters et al., 2009; Wong et al., 2013), our work revealed a reliable relationship between fingertip size and (edge-orientation acuity during active movement. This relationship likely is thought to reflect the increased innervation density of Merkel and Meissner mechanoreceptors in smaller fingertips (Peters et al., 2009). We believe our finding is rather strong support for this mechanistic explanation because, in our paradigm, participants could have changed the way they contacted the edges or moved their finger and perhaps compensate for such effects. However, we found no reliable difference in these parameters, either as a discrete function of gender or as it related with fingertip size. Of course our results are not definitive because, as previously elaborated in detail (Peters et al., 2009), there exist a range of factors that correlate with fingertip size that may actually underlie the effect being described.

## Acknowledgements

This work was supported by the Canadian Institutes of Health Research (Foundation Grant to JAP: 353197) and the Government of Ontario (Early Researcher Award to JAP). JAP received a salary award from the Canada Research Chairs Program.

## References

Bensmaia, S.J., Hsiao, S.S., Denchev, P.V., Killebrew, J.H., and Craig, J.C. (2008). The tactile perception of stimulus orientation. Somatosens. Mot. Res. 25, 49–59.

Chapman C.E. (1994). Active versus passive touch: factors influencing the transmission of somatosensory signals to primary somatosensory cortex. Can. J. Physiol. Pharmacol. 72, 558–570.

Gibson J.J. (1962). Observations on active touch. Psychol. Rev. 69, 477–491.

Goldreich, D., and Kanics, I.M. (2003). Tactile acuity is enhanced in blindness. J. Neurosci. Off. J. Soc. Neurosci. 23, 3439–3445.

Goldreich, D., and Kanics, I.M. (2006). Performance of blind and sighted humans on a tactile grating detection task. Percept. Psychophys. 68, 1363–1371.

Heller M.A. (1989). Texture perception in sighted and blind observers. Percept. Psychophys. 45, 49–54.

Johansson, R.S., and Flanagan, J.R. (2009). Coding and use of tactile signals from the fingertips in object manipulation tasks. Nat. Rev. Neurosci. 10, 345–359.

Lamb G.D. (1983). Tactile discrimination of textured surfaces: psychophysical performance measurements in humans. J. Physiol. 338, 551–565.

Lechelt E.C. (1992). Tactile spatial anisotropy with static stimulation. Bull. Psychon. Soc. 30, 140–142.

Lederman S.J. (1981). The perception of surface roughness by active and passive touch. Bull. Psychon. Soc. 18, 253–255.

Lederman, S.J., and Klatzky, R.L. (1987). Hand movements: a window into haptic object recognition. Cognit. Psychol. 19, 342–368.

Peters, R.M., Hackeman, E., and Goldreich, D. (2009). Diminutive digits discern delicate details: fingertip size and the sex difference in tactile spatial acuity. J. Neurosci. Off. J. Soc. Neurosci. 29, 15756–15761.

Peters, R.M., Staibano, P., and Goldreich, D. (2015). Tactile orientation perception: an ideal observer analysis of human psychophysical performance in relation to macaque area 3b receptive fields. J. Neurophysiol. 114, 3076–3096.

Pruszynski, J.A., and Johansson, R.S. (2014). Edge-orientation processing in first-order tactile neurons. Nat. Neurosci. 17, 1404–1409.

Pruszynski, J.A., Johansson, R.S., and Flanagan, J.R. (2016). A Rapid Tactile-Motor Reflex Automatically Guides Reaching toward Handheld Objects. Curr. Biol. 26, 788–792.

Pruszynski, J.A., Flanagan, J.R., and Johansson, R.S. (2018). Fast and accurate edge orientation processing during object manipulation. ELife 7.

Van Boven, R.W., Hamilton, R.H., Kauffman, T., Keenan, J.P., and Pascual-Leone, A. (2000). Tactile spatial resolution in blind braille readers. Neurology 54, 2230–2236.

Vega-Bermudez, F., Johnson, K.O., and Hsiao, S.S. (1991). Human tactile pattern recognition: active versus passive touch, velocity effects, and patterns of confusion. J Neurophysiol 65, 531–546.

Verrillo, R.T., Bolanowski, S.J., and McGlone, F.P. (1999). Subjective magnitude of tactile roughness. Somatosens. Mot. Res. 16, 352–360.

Wong, M., Peters, R.M., and Goldreich, D. (2013). A physical constraint on perceptual learning: tactile spatial acuity improves with training to a limit set by finger size. J. Neurosci. Off. J. Soc. Neurosci. 33, 9345–9352.

